# Novel regulation of the transcription factor ZHX2 by N-terminal methylation

**DOI:** 10.1101/2021.10.22.465472

**Authors:** Meghan M. Conner, Haley V. Parker, Daniela R. Falcone, Gehoon Chung, Christine E. Schaner Tooley

## Abstract

N-terminal methylation (Nα-methylation) by the methyltransferase NRMT1 is an important post-translational modification that regulates protein-DNA interactions. Accordingly, its loss impairs functions that are reliant on such interactions, including DNA repair and transcriptional regulation. Global loss of Nα-methylation results in severe developmental and premature aging phenotypes, but given over 300 predicted substrates, it is hard to discern which physiological substrates contribute to each phenotype. One of the most striking phenotypes in NRMT1 knockout (*Nrmt1^-/-^*) mice is early liver degeneration. To identify the disrupted signaling pathways leading to this phenotype and the NRMT1 substrates involved, we performed RNA-sequencing analysis of control and *Nrmt1^-/-^* adult mouse livers. We found both a significant upregulation of transcripts in the cytochrome P450 (CYP) family and downregulation of transcripts in the major urinary protein (MUP) family. Interestingly, transcription of both families is inversely regulated by the transcription factor zinc fingers and homeoboxes 2 (ZHX2). ZHX2 contains a non-canonical NRMT1 consensus sequence, indicating its function could be directly regulated by Nα-methylation. We confirmed misregulation of CYP and MUP mRNA and protein levels in *Nrmt1^-/-^* livers and verified NRMT1 can methylate ZHX2 *in vitro*. In addition, we used mutants of ZHX2 that cannot be methylated to directly demonstrate Nα-methylation promotes ZHX2 transcription factor activity. Finally, we show *Nrmt1^-/-^* mice also exhibit early postnatal de-repression of ZHX2 targets involved in fetal liver development. Taken together, these data implicate continual ZHX2 misregulation as a driving force behind the liver phenotype seen in *Nrmt1^-/-^* mice.

## Introduction

N-terminal methylation (Nα-methylation), occurring on the α-amino group at the N-terminus of proteins, is a post-translation modification (PTM) that can affect protein stability and nucleic acid-binding activity (1, 2). Nα-methylation was first observed as a PTM many decades ago (3), but it was only ten years ago that the responsible methyltransferase was identified as <u>N</u>-terminal <u>R</u>CC1 <u>m</u>ethyl<u>t</u>ransferase 1 (NRMT1) (4). NRMT1 is the major eukaryotic Nα-trimethylase that sequentially mono-, di-, and trimethylates protein N-termini after removal of the initiating methionine (Met) (4). It was originally thought that NRMT1 could only act on N-termini with the canonical consensus sequence X-Pro-Lys, where X is Ala, Pro, or Ser (3). However, comprehensive sequence analysis identified a greatly expanded non-canonical consensus sequence which allows A/P/S/G/M in the first position, A/P/S/G/M/E/N/Q in the second position, and K/R in the third position after Met cleavage (5). This expanded consensus sequence predicts over 300 potential NRMT1 targets, with dozens being verified in recent years (5–11).

The most well-studied function of Nα-methylation by NRMT1 is its ability to promote DNA-binding of its substrates (2, 4, 6, 8, 10). The first characterized substrate of NRMT1, regulator of chromatin condensation 1 (RCC1), requires Nα-methylation for association with chromatin and subsequent proper spindle assembly and chromosome segregation during mitosis (2, 4). Use of a methyl-deficient mutant with Glu replacing Lys in the third position of the RCC1 N-terminal sequence (K4Q mutant) revealed RCC1 does not co-localize with chromatin when not methylated (2), and knockdown of NRMT1 recapitulates this phenotype (4). *In vitro* binding assays confirmed increased association of methylated RCC1 to DNA and not histone proteins (2). RCC1 was the first protein shown to require Nα-trimethylation by NRMT1 for proper association with DNA, but several other similarly regulated proteins have now been identified, including the centromere protein B (CENP-B) (6), damaged DNA-binding protein 2 (DDB2) (8), and myosin light chain 9 (MYL9) (10).

CENP-B, which has a GPK canonical consensus, binds to a CENP-B box sequence in centromeric DNA, and a K4Q CENP-B mutant was unable to bind the CENP-B box as strongly as wild-type (WT) (6). WT DDB2, which has an APK canonical consensus, was recruited to DNA damage foci at a greater rate than the K4Q unmethylatable DDB2 mutant (8), and a K4Q mutant of the transcription factor MYL9 decreased its occupancy at target promoters (10). The N-terminal Ser-Ser-Lys (SSK) consensus sequence of MYL9 makes it the first non-canonical target identified whose DNA binding is regulated by Nα-methylation (10). Interestingly, the SSK consensus also allows for Nα-acetylation of MYL9, and while Nα-methylation regulates its nuclear role as a transcription factor, we have also shown that Nα-acetylation regulates its cytoplasmic roles in cell migration (10).

To more comprehensively identify additional targets of NRMT1 regulation, we performed RNA-sequencing (RNA-seq) analysis on the livers of *Nrmt1^-/-^* mice. Livers were chosen because one of the most striking phenotypes of the *Nrmt1^-/-^* mice is the early degeneration of this tissue (12). When compared to transcript levels in WT mice livers, hundreds of differentially expressed genes were identified in *Nrmt1^-/-^* mice. Most notably, transcripts belonging to the Cytochrome p450 (CYP) family were more highly expressed in *Nrmt1^-/-^* livers, and transcripts of the major urinary protein (MUP) family showed reduced expression in *Nrmt1^-/-^* livers. Interestingly, these families are inversely regulated by a common transcription factor, zinc fingers and homeoboxes 2 (ZHX2). ZHX2 both represses expression of many CYP genes and activates expression of many MUP genes (13, 14).

ZHX2 is a member of a family of transcriptional factors that also includes ZHX1 and ZHX3, all of which can homo- or heterodimerize (15, 16). The structure of ZHX2, including two zinc finger motifs and five homeodomains, suggests it interacts with both other proteins and nucleic acids (17, 18). ZHX2 has been implicated in liver development, through its postnatal repression of fetal developmental regulators alphafetoprotein (AFP), glypican-3 (GPC3), and lncRNA H19 (19, 20). It has also been implicated in numerous diseases, including liver and renal cancers (21–24). In hepatocellular carcinoma (HCC), ZHX2 acts as a tumor suppressor and its loss from the nucleus leads to increased proliferation through increased expression of its targets cyclin A and cyclin E (22). However, in clear cell renal cell carcinoma (ccRCC), ZHX2 acts as an oncogene by activating the nuclear factor kappa B (NF-κB) pathway, and its loss slows cell growth (24).

Given our RNA-seq data and the fact that ZHX2 has a non-canonical Ala-Ser-Lys (ASK) NRMT1 consensus sequence, we decided to test whether ZHX2 is a direct target of NRMT1 and if Nα-methylation of ZHX2 regulates its function as a transcription factor. Here we verify that NRMT1 can methylate ZHX2 *in vitro*, and through the use of a K4Q unmethylatable ZHX2 mutant, we also find that Nα-methylation of ZHX2 is required for activation of its target genes. Additionally, we show that while *Nrmt1^-/-^* mice postnatally repress *Afp* expression similar to WT mice, expression of both *Gpc3* and *H19* is de-repressed by 14 days after birth. Taken together, we find that ZHX2 is a novel target of NRMT1 and loss of Nα-methylation leads to aberrant regulation of ZHX2 function as a transcription factor, resulting in altered liver metabolic and developmental pathways.

## Materials and methods

### RNA sequencing

RNA was isolated from the livers of three 3-month-old male control C57BL/6J and *Nrmt1^-/-^* mice using TRIzol reagent (Thermo Fisher Scientific, Waltham, MA) as previously described (12). Two mRNA libraries per sample were made per the standard Illumina protocol and sequenced on the Illumina NextSeq 500 System at the University of Louisville Center for Genetics and Molecular Medicine. Eleven of the twelve libraries produced analyzable data, so all but one of the control samples are represented as duplicates. There were 23,545 genes represented in the 11 samples. Genes with low expression were eliminated and those represented by at least 1 count per 1 million reads in at least 3 samples were included in subsequent analysis. Across the 11 samples, 12,612 genes were retained. The trimmed mean of M-values (TMM) method (25) implemented in edgeR (3.4) was used to calculate normalization factors and examine inter-sample reproducibility. The reads were mapped to the Ensembl *Mus musculus* genome provided by Illumina and STAR (2.5.3a) aligner using default settings. The dispersion parameter for each gene was estimated using the quantile-adjusted conditional maximum likelihood (qCML) method, appropriated for experiments with a single factor. The functions estimateCommonDisp and estimateTagwiseDisp were used to estimate dispersion. Differential expression was tested using the exact test based on qCML methods. The Benjamini-Hochberg correction was used with a false discovery cut-off of 0.1. With these parameters, 102 genes (0.809%) were shown positively-regulated in control vs. *Nrmt1^-/-^* mice and 119 genes (0.944%) were negatively-regulated.

### Gene ontology (GO) analysis

Gene ontology (GO) analysis of the top 50 positively-regulated and negatively-regulated transcripts in the *Nrmt1^-/-^* mice and control mice was performed using the Database for Annotation, Visualization and Integrated Discovery (DAVID) (ver 6.8). Genes were grouped by both molecular function (MF) and biological process (BP).

### Molecular cloning

Full-length wild-type (WT) or methylation-deficient mutant K4Q (ASQ) ZHX2 were amplified from human ZHX2 cDNA in pOTB7 (Horizon Discovery, Waterbeach, United Kingdom) and cloned into the XbaI and BamHI sites of pKGFP2 (a generous gift from Dr. Ian Macara) to put a C-terminal GFP tag on ZHX2. The 5’ primer for the WT construct was 5’-TCTAGAATGGCTAGCAAACGAAAATC-3’and the 5’ primer for the ASQ construct was 5’-TCTAGAATGGCTAGCCAACGAAAATC-3’. The 3’ primer for both was 5’-GGATCCGGCCTGGCCAGCCTCTGCAG-3’. To make an expression vector that simultaneously produces NRMT1 and NRMT2, each were cloned into pETDuet-1 (EMD Millipore, Burlington, MA). First, the human NRMT1 ORF (4) was amplified to introduce a 5’ HindIII restriction site and a 3’NotI restriction site with a stop codon and subcloned into pETDuet-1 using HindIII and NotI to put an N-terminal His tag on NRMT1. Next, the human NRMT2 ORF (26) was amplified to introduce a 5’ NdeI restriction site and a 3’ BglII restriction site with a stop codon. As NRMT1 naturally contains a BglII restriction site, Quikchange site-directed mutagenesis (Agilent, Santa Clara, CA) was used on the NRMT1-pETDuet-1 vector to introduce a silent mutation at Ile214 (ATC to ATA) and disrupt the BglII site. The amplified NRMT2 ORF was then subcloned into this NRMT1-pETDuet-1 vector using NdeI and BglII, which will result in the production of untagged NRMT2. The following primers were used: 5HindIIINRMT1: 5’-GCAAGCTTACGAGCGAGGTGATAGAAGAC-3’; 3NotIStopNRMT1: 5’-CCGCGGCCGCTCATCTCAGGGCAAAGCTATA-3’; hsNRMT1BglIIQC For: 5’-GGAGAACCTCCCCGATGAGATATACCATGTCTATAGC-3’; hsNRMT1BglIIQC Rev: 5’-GCTATAGACATGGTATATCTCATCGGGGAGGTTCTCC-3’; 5NdeINRMT2: 5’-GGCATATGGCCCACCGGGGAGCCCA-3’; 3BglIINRMT2Stop: 5’-GCAGATCTTCAGGAGTGTCTGTCGCTGTG-3’. All constructs generated were verified by DNA sequencing.

### Cell culture, lipofectamine transfection, and proliferation assays

Human renal carcinoma 786-O cells (ATCC, Manassas, VA) were cultured in RPMI 1640 media (Corning, Corning, NY) supplemented with 10% fetal bovine serum (FBS; Atlanta Biologicals, Atlanta, GA) and 1% penicillin-streptomycin (P/S; Thermo Fisher). Cells were grown at 37°C and 5% CO_2_ on tissue culture treated plastic (Corning). Human 786-O cells were transfected with 1.5 μg WT-ZHX2-GFP or K4Q-ZHX2-GFP using Lipofectamine 2000 reagent (Thermo Fisher) per manufacturer instructions. After 24 hours, GFP expression was confirmed with an EVOS fluorescent microscope (Thermo Fisher) and cells were harvested for qRT-PCR analysis. Lentivirus was generated through co-transfection of HEK293T cells with 50 μg pGIPZ vector containing shRNAs against human or mouse NRMT1 (Horizon Discovery), 37.5 μg psPAX2 packaging vector, and 15 μg pMD2.G envelope plasmid using calcium phosphate transfection as previously described (27). Transduction of 786-O cells with each lentivirus was performed at a multiplicity of infection (MOI) of 5. Cells transduced with lentivirus expressing the shRNA against mouse NRMT1 were used as the control. Cell proliferation experiments were carried out using the CellTiter 96 Aq_ueous_One Solution Cell Proliferation Assay reagent (Promega, Madison, WI). For these experiments, 1000 786-O cells were plated in triplicate in a 96-well plate. Daily cell number was determined by addition of 20 μl of Aq_ueous_One Solution and measurement of absorbance at 490 nm using a Cytation 5 imaging system (BioTek, Winooski, VT). Fold change was calculated by dividing by absorbance measurements taken at day zero.

### Western blots

Liver lysates were generated by homogenizing livers into a buffer containing 15 mM Tris (pH 7.6), 0.25 M sucrose, 2 mM EDTA, 1 mM EGTA, 10 mM Na3VO4, 25 mM NaF, 10 mM sodium pyrophosphate, 1 mM PMSF, 1 μg/mL leupeptin, and 1 μg/mL aprotinin. Protein samples were quantified using a Bradford assay and an equal concentration was loaded for each sample in 5 μl of Laemmli Sample Buffer (60 mM Tris (pH 6.8), 2% SDS, 10% glycerol, 5% β-mercaptoethanol, 0.01% bromophenol blue). Samples were separated on 10% SDS-PAGE gels and transferred to nitrocellulose membranes (Bio-Rad, Hercules, CA) using a Trans-Blot Turbo Transfer System (Bio-Rad). Membranes were incubated for one hour at room temperature in blocking buffer (5% w/v non-fat dry milk in TBS + 0.1% Tween 20 (TBS-T)). Primary and secondary antibodies were incubated in 5% w/v non-fat dry milk in TBS-T according to manufacturer’s instructions. Dilutions used for primary antibodies were rabbit anti-GAPDH (1:1000; 2275-PC-100, Trevigen, Gaithersburg, MD), rabbit anti-β-tubulin (9F3) (1:1000; Cell Signaling Technology, Danvers, MA), rabbit anti-NRMT1 (1:1000; (4)), mouse anti-ZHX2 (D-2) (1:1000; sc-393399, Santa Cruz Antibodies, Dallas, TX), rabbit anti-CYP26A1 (1:1000; PA5-28165, Invitrogen, Waltham, MA), and rabbit anti-MUP (1:1000; PA5-112879, Invitrogen). Secondary antibodies used were donkey anti-mouse or donkey anti-rabbit (1:5000; Jackson ImmunoResearch, West Grove, PA). Blots were developed on a ChemiDoc Touch imaging system (Bio-Rad) using Clarity Western ECL Substrate (Bio-Rad) or SuperSignal West Femto Maximum Sensitivity Substrate (Thermo Fisher). Quantification of blots was performed using ImageJ 1.53e (NIH, Bethesda, MD).

### Real time PCR analysis

Mouse livers or 786-O pellets were homogenized in TRIzol (Invitrogen) and mixed with chloroform to extract RNA. RNA was pelleted in isopropanol, washed with ethanol, and resuspended in 40 - 100 μl sterile water. cDNA was synthesized from 1 μg RNA using the SuperScript First-Strand Synthesis System (Invitrogen) and diluted 1:10 in sterile water. For quantitative RT-PCR, 2 μl of each sample was used with 2X SYBR green Master Mix and the CFX96 Touch Real-Time PCR Detection System (Bio-Rad). Samples were run in triplicate. Transcript expression levels were determined using the ΔΔCT quantification method. Primer sequences (Invitrogen) were as follows: mouse *β-tubulin* forward 5’-TGTCCTGGACAGGATTCGC-3’ and reverse 5’-CTCCATCAGCAGGGAGGTG-3’; mouse *Cyp17a1* forward 5’-CAATGACCGGACTCACCTCC-3’ and reverse 5’-GATCACTGTGTGTCCTTCG-3’; mouse *Cyp2a4* forward 5’-CGTTGGGAACTTCCTTCAGC-3’ and reverse 5’-CTTCCTTGACTGTCTCCTGT-3’; mouse *Mup1* forward 5’-CACACTACAGATCGCTGCCT-3’and reverse 5’-CAGGATGGAATGCAGATCAC-3’; mouse *Mup7* forward 5’-TACCAAAATGAAGATGCTGC-3’and reverse 5’-ATAGTATGCCATTCCCCA-3’; human *GAPDH* forward 5’-ACAGCCTCAAGATCATCAGCAA-3’ and reverse 5’-CCATCACGCCACAGTTTCC-3’; human *BCL2* forward 5’-ATCGCCCTGTGGATGACTGAGT-3’; human *ICAM1* forward 5’-CTCAGTTTCCCAGCGACAGG-3’ and reverse 5’-GGAAGCTGCGTGATCCCTAC-3’; human *IL8* forward 5’-TCTGACATAATGAAAAGATGAGGGT-3’ and reverse 5’-CCTTCCGGTGGTTTCTTCCT-3’ (24); mouse *Afp* forward 5’-ATCAGTGTCTGCTGGCACGCA-3’ and reverse 5’-GGCTGGGGCATACATGAAGGGG-3’ (13); mouse *H19* forward 5’-GTGTCACCAGAAGGGGAGTG-3’ and reverse 5’-AGTGCCTCATGGGAATGGTG-3’; and mouse *Gpc3* forward 5’-CCAGGTTTCCAAGTCACTG-3’ and reverse 5’-CTTGAGGTGGTCGGTAGTGT-3’ (28).

### In vitro methylation assays

Recombinant His_6_-tagged NRMT1 or His_6_-tagged NRMT1+NRMT2 proteins were purified as previously described (4, 29). Briefly, pET15b-NRMT1 (5) or pETDuet-1-NRMT1/NRMT2 was expressed in BL21 Star (DE3) *E. coli* (Thermo Fisher) and purified on Ni^2+^-NTA beads (Qiagen, Hilden, Germany). NRMT1 and NRMT2 complex together (1), so the His-tagged NRMT1 expressed by the pET15b will also pull out the untagged NRMT2. The methyltransferase assays were conducted using the MTase-Glo Methyltransferase Assay (Promega) following the protocol from the manufacturers. Briefly, 0.2 μM NRMT1 or 0.2 μM NRMT1+NRMT2 was incubated in wells on a 96-well plate with 40 μM of synthetic peptide substrate corresponding to the first 15 N-terminal amino acids of ZHX2 (minus the initiating methionine) (AnaSpec, Fremont, CA) in the presence of 40 μM S-adenosyl methionine (SAM). Reactions were incubated at room temperature and stopped after 20 minutes with the addition of 0.5% trifluoroacetic acid. The MTase-Glo detection reagents were added according to the manufacturers protocols, and luminescence was measured using the Cytation5 Imaging System (BioTek). Background signal was measured for each condition through the inclusion of no substrate control reactions and was subtracted from respective experimental reactions.

### Statistical analyses

All statistical analysis was performed using Prism 6 software (GraphPad, San Diego, CA). Statistical test used is denoted in the figure legends. Results shown are mean +/- standard deviation or standard error of the mean (SEM), as denoted in the figure legends.

## Results

### RNA sequencing of Nrmt1^-/-^ livers

Our initial characterization of the NRMT1 knockout (*Nrmt1^-/-^*) mouse was striking, given the widespread aging phenotypes we observed including extensive liver degeneration (12). To begin to understand what disrupted signaling pathways lead to this degeneration and what NRMT1 substrates are involved, we performed RNA sequencing (RNA-seq) analysis on livers from 3-month-old *Nrmt1^-/-^* and control wild type (WT) C57BL/6J male mice. The RNA-seq revealed over two hundred transcripts with differential expression between the two groups (102 genes positively-regulated in WT, 119 genes negatively-regulated).

Analysis of the top 50 up- and down-regulated genes showed expected misregulation of genes involved in liver metabolism, small molecule and pheromone processing, and transport (Figures 1 and 2, Supplemental Tables 1 and 2). *Nrmt1* was one of the top ten transcripts with higher expression in WT livers, further validating the sequencing results (Figure 1(a)). Interestingly, several genes belonging to the major urinary protein (MUP) family were additionally upregulated in WT/down-regulated in the *Nrmt1^-/-^* livers, including *Mup8, Mup14, Mup7, Mup15, Mup1, Mup12*, and *Mup21* (Figure 1(a)). MUP proteins are secreted by the liver, bind pheromones and other lipophilic molecules, regulate their transportation, and are ultimately secreted in the urine (30). Gene ontology (GO) analysis of the top 50 genes down-regulated in *Nrmt1^-/-^* livers confirms enrichment of MUP transcripts in the dataset, showing the highest functional enrichment in genes involved in transporter activity and small molecule and pheromone binding (Figure 1(b)). Enriched biological processes include ion and transmembrane transport, as well as negative regulation of lipid biosynthetic processes (Figure 1(c)).

**Figure 1.**
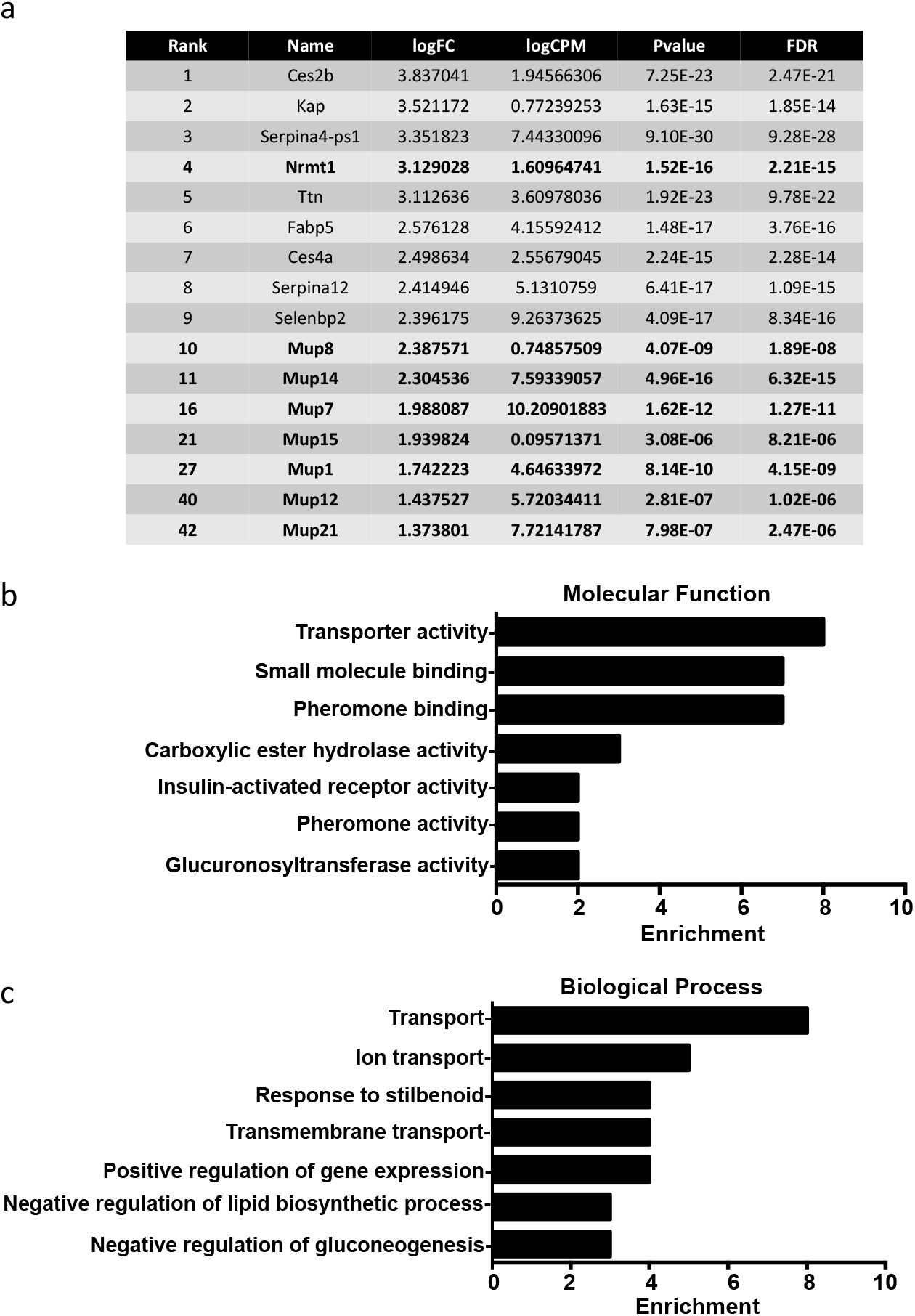
Transcripts down-regulated in *Nrmt1^-/-^* mice. (a) The top ten transcripts with lower expression in *Nrmt1^-/-^* mice include *Nrmt1* and *Mup8*. Extending past the top ten into the top 50 reveals down-regulation of several other major urinary protein (*Mup*) transcripts. Gene ontology analysis provided enriched (b) molecular functions and (c) biological processes of the top 50 genes with down-regulated expression.

**Figure 2.**
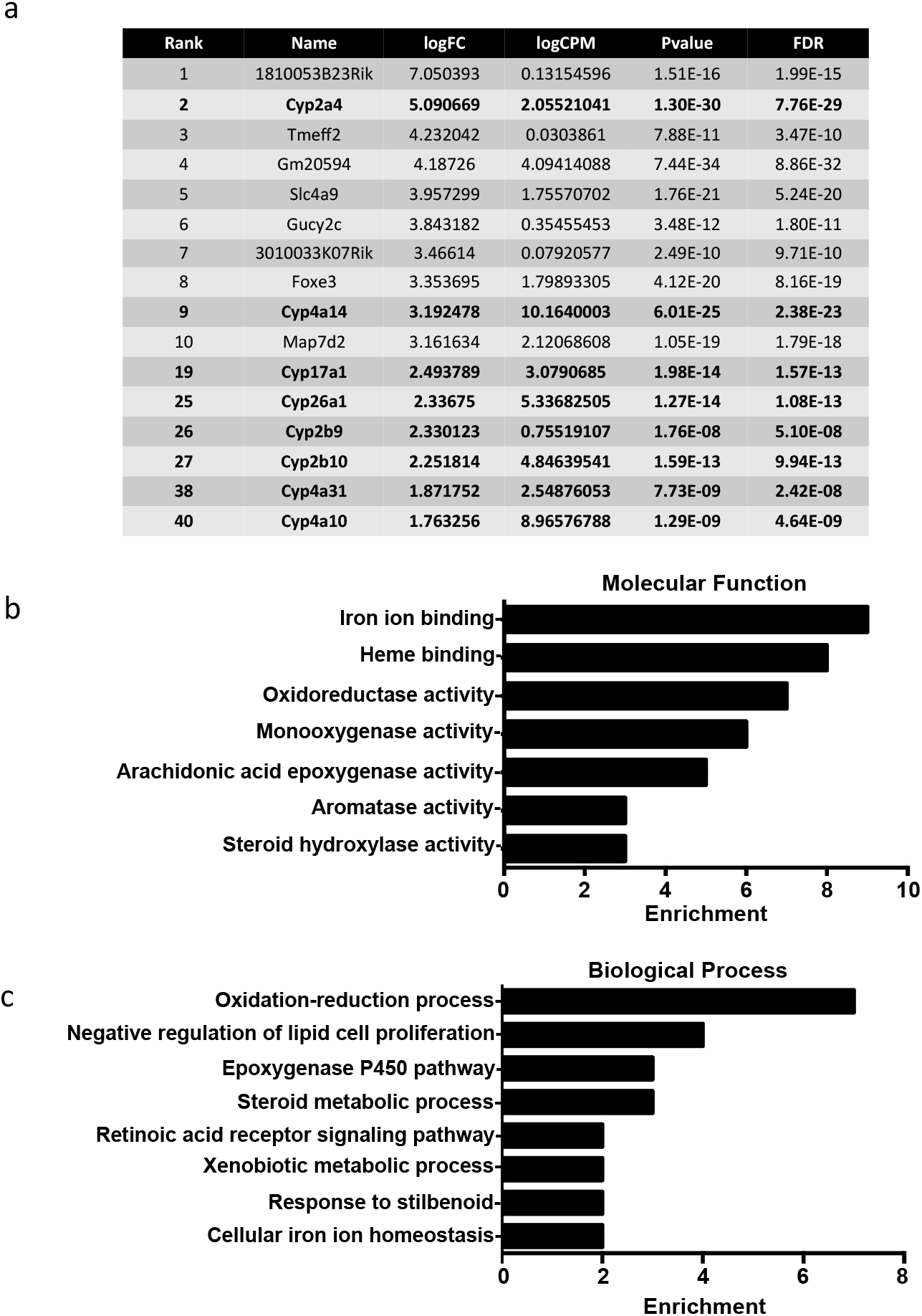
Transcripts upregulated in *Nrmt1^-/-^* mice. (a) The top ten transcripts with higher expression in *Nrmt1^-/-^* mice include *Cyp2a4* and *Cyp4a14*. Extending past the top ten into the top 50 reveals upregulation of several other cytochrome p450 (*Cyp*) transcripts. Gene ontology analysis provided enriched (b) molecular functions and (c) biological processes of the top 50 genes with upregulated expression.

Conversely, several genes belonging to the cytochrome p450 (CYP) family showed down-regulated expression in WT livers/upregulation in *Nrmt1^-/-^* livers, including *Cyp2a4, Cyp4a14, Cyp26a1, Cyp17a1, Cyp2b10, Cyp4a10, Cyp4a31*, and *Cyp2b9* (Figure 2(a)). The CYP family is comprised of hemeproteins that regulate steroid hormone synthesis and the metabolism of drugs and other xenobiotics, primarily through their monooxygenase hydroxylation activity (31). GO analysis of the top 50 genes upregulated in *Nrmt1^-/-^* livers confirms enrichment of CYP genes in the dataset, showing the highest functional enrichment in genes involved in iron ion binding, heme binding, and oxidoreductase and monooxygenase activity (Figure 2(b)). Also enriched are epoxygenase, aromatase, and steroid hydroxylase activity, all functions of CYP enzymes (32–34). Enriched biological processes include oxidation-reduction processes, epoxygenase P450 pathway, steroid metabolic processes, and xenobiotic metabolic processes (Figure 2(c)).

### Verification of RNA sequencing

To verify the RNA-seq data at the mRNA level, quantitative real-time PCR (qRT-PCR) was performed. First, as a control, *Nrmt1* mRNA levels were compared between WT and *Nrmt1^-/-^* livers and found to be significantly higher in WT samples (Figure 3(a)). qRT-PCR analysis of two representative *Mup* genes, *Mup1* and *Mup7*, also showed significantly higher levels in WT samples (Figure 3(b)). *Mup1* and *Mup7* transcripts were chosen for analysis due to their high differential expression between WT and *Nrmt1^-/-^* livers and regions of sequence disparity between other *Mup* genes that allowed for specific primer design. qRT-PCR analysis of two representative *Cyp* genes from the RNA-seq data, *Cyp17a1* and *Cyp2a4*, also verified the RNA-seq data, showing significantly lower expression in WT livers (Figure 3(c)). These Cyp genes were also chosen for their high differential expression in the RNA-seq data and ease of primer design.

**Figure 3.**
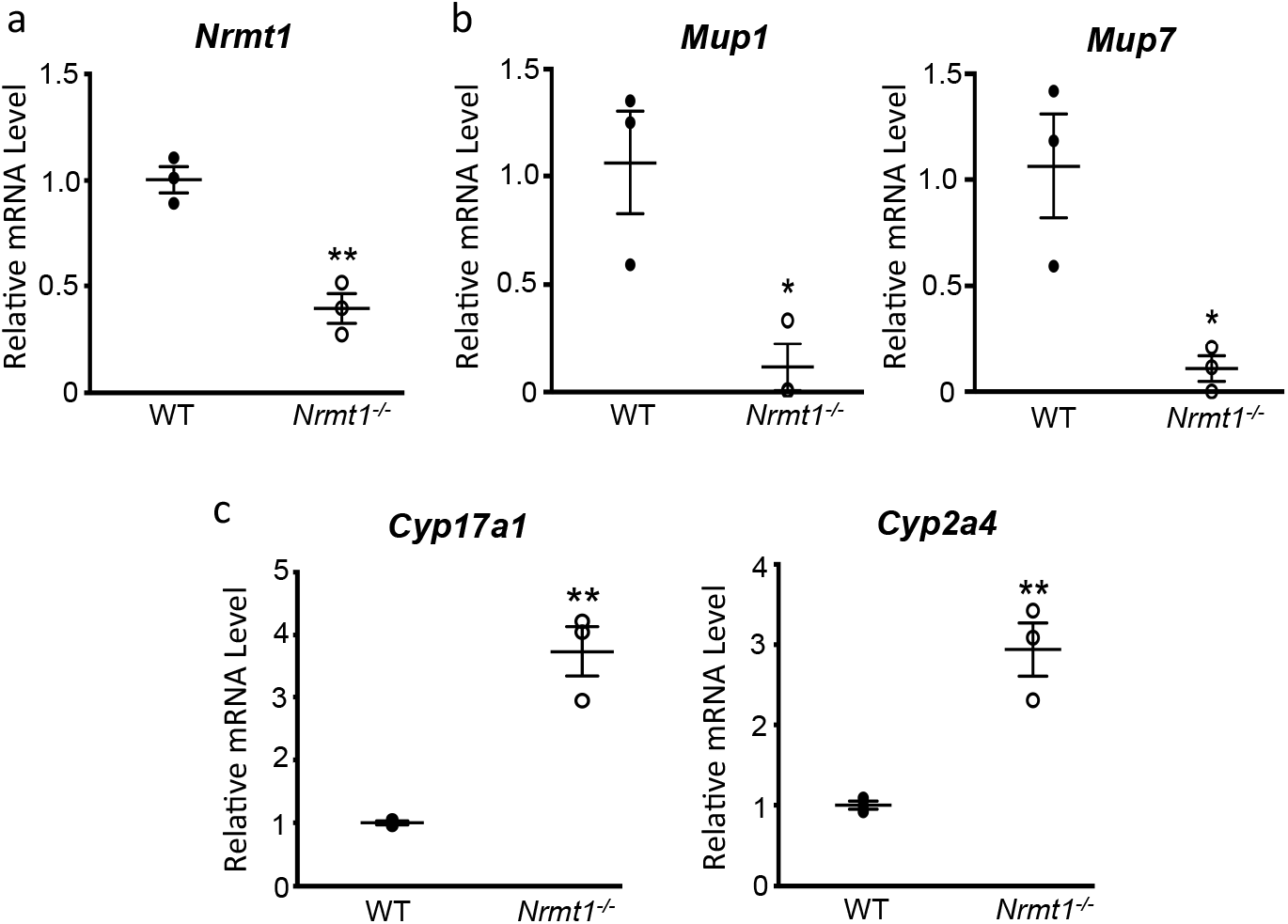
qRT-PCR verification of RNA sequencing. (a-c) qRT-PCR analysis of wildtype (WT) and Nrmt1 knockout (*Nrmt1^-/-^*) mice shows (a) decreased *Nrmt1* expression, (b) decreased *Mup1* and *Mup7* expression, and (c) increased *Cyp17a1* and *Cyp2a4* expression. * denotes p < 0.05 and ** denotes p < 0.01 as determined by unpaired t-test. n = 3. Error bars represent ± standard error of the mean (SEM).

To verify the RNA-seq data at the protein level, protein expression changes between WT and *Nrmt1^-/-^* livers were measured by western blot. Again, verifying the RNA-seq data, MUP protein expression was significantly higher in the WT liver samples (Figure 4(a,b)). Total MUP protein was analyzed by western blot, as opposed to individual MUP proteins, as the MUP proteins are highly similar and specific antibodies were not readily available. As at least seven *Mup* genes are downregulated in *Nrmt1^-/-^* livers, we reasoned correctly this should result in a significant reduction in total MUP protein. We also confirmed that Cyp26a1 protein levels were significantly lower in WT livers as compared to *Nrmt1^-/-^* livers (Figure 4(a,c)). Cyp26a1 was chosen for its high differential expression and antibody quality.

**Figure 4.**
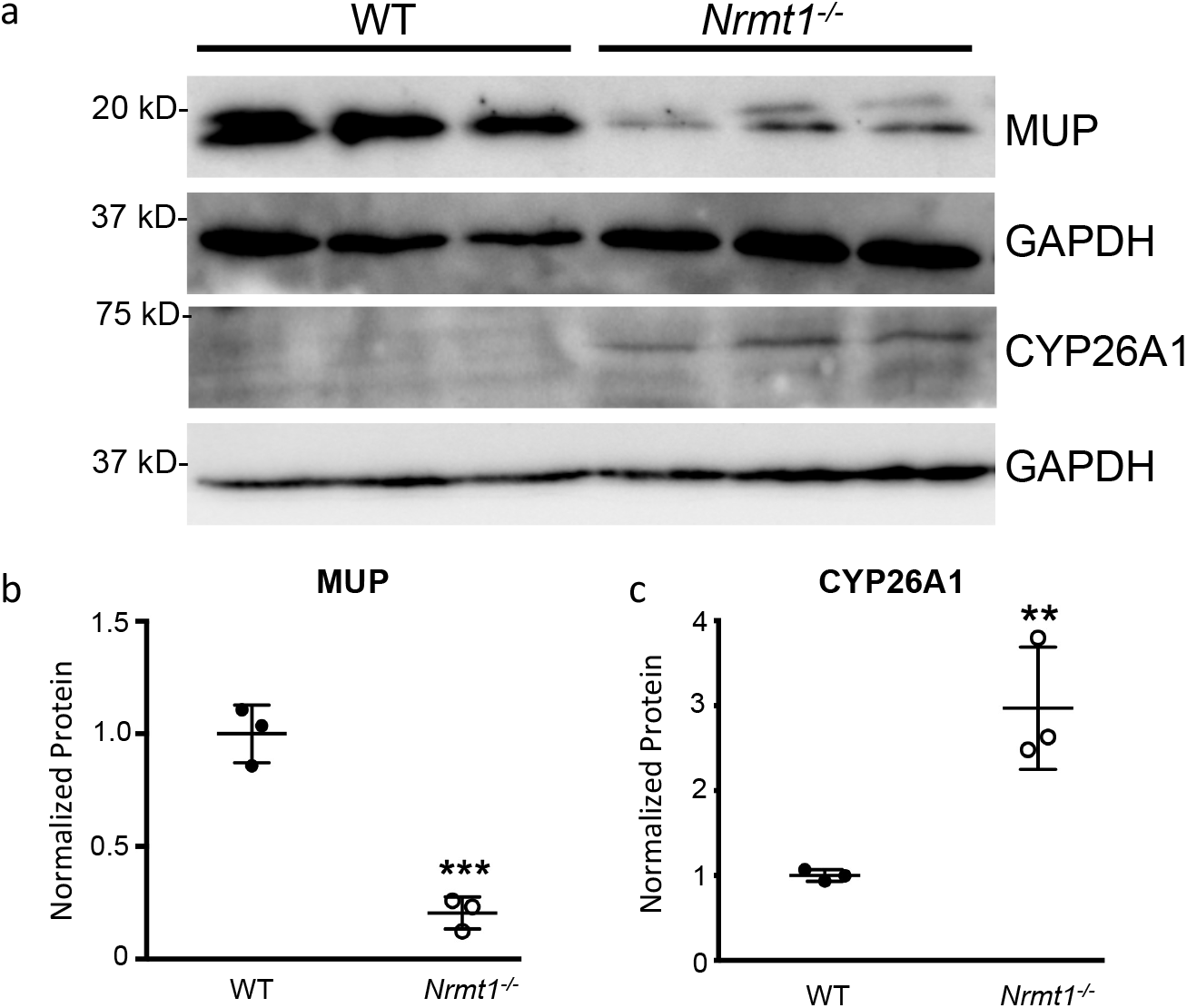
Western blot verification of RNA sequencing. (a) Western blot analysis of wild-type (WT) and Nrmt1 knockout (*Nrmt1^-/-^*) mouse livers verifies (b) significant downregulation of MUP protein levels in *Nrmt1^-/-^* mice and (c) significant upregulation of CYP26A1 protein levels in *Nrmt1^-/-^* mice. GAPDH is used as a loading control. * p < 0.01 and *** denotes p < 0.001 as determined by unpaired t-test. n = 3. Error bars represent ± standard error of the mean (SEM).

### ZHX2 function is regulated by Nα-methylation

The discovery of both down-regulation of *Mup* genes and upregulation of *Cyp* genes in *Nrmt1^-/-^* livers was interesting, as both are regulated inversely by the transcription factor zinc fingers and homeoboxes 2 (ZHX2) (13, 14). ZHX2 has been shown to both promote *Mup* expression (13) and repress *Cyp* expression (14). Given the RNA-seq results show lowered Mup expression and increased Cyp expression in *Nrmt1^-/-^* livers, they indicate ZHX2 is unable to function optimally as a transcription factor in the absence of NRMT1.

Nα-methylation by NRMT1 has been shown to regulate protein/DNA interactions, and its loss has been shown to disrupt processes that rely on these interactions (2, 6, 8, 10). Analysis of the N-terminal sequence of ZHX2 showed it has a non-canonical NRMT1 consensus sequence, Ala-Ser-Lys (ASK – after initiating methionine cleavage), indicating ZHX2 could be a direct target of NRMT1. To verify this, *in vitro* methylation assays were performed with an N-terminal ZHX2 peptide and recombinant human NRMT1 alone or in combination with recombinant human NRMT2. We have previously shown that non-canonical substrates can be methylated by NRMT1, but this activity is significantly increased by complex formation between NRMT1 and NRMT2 (1). NRMT1 alone was able to methylate the ZHX2 peptide, and addition of NRMT2 significantly increased this methylation, confirming that ZHX2 is a non-canonical target of NRMT1 (Figure 5(a)).

**Figure 5.**
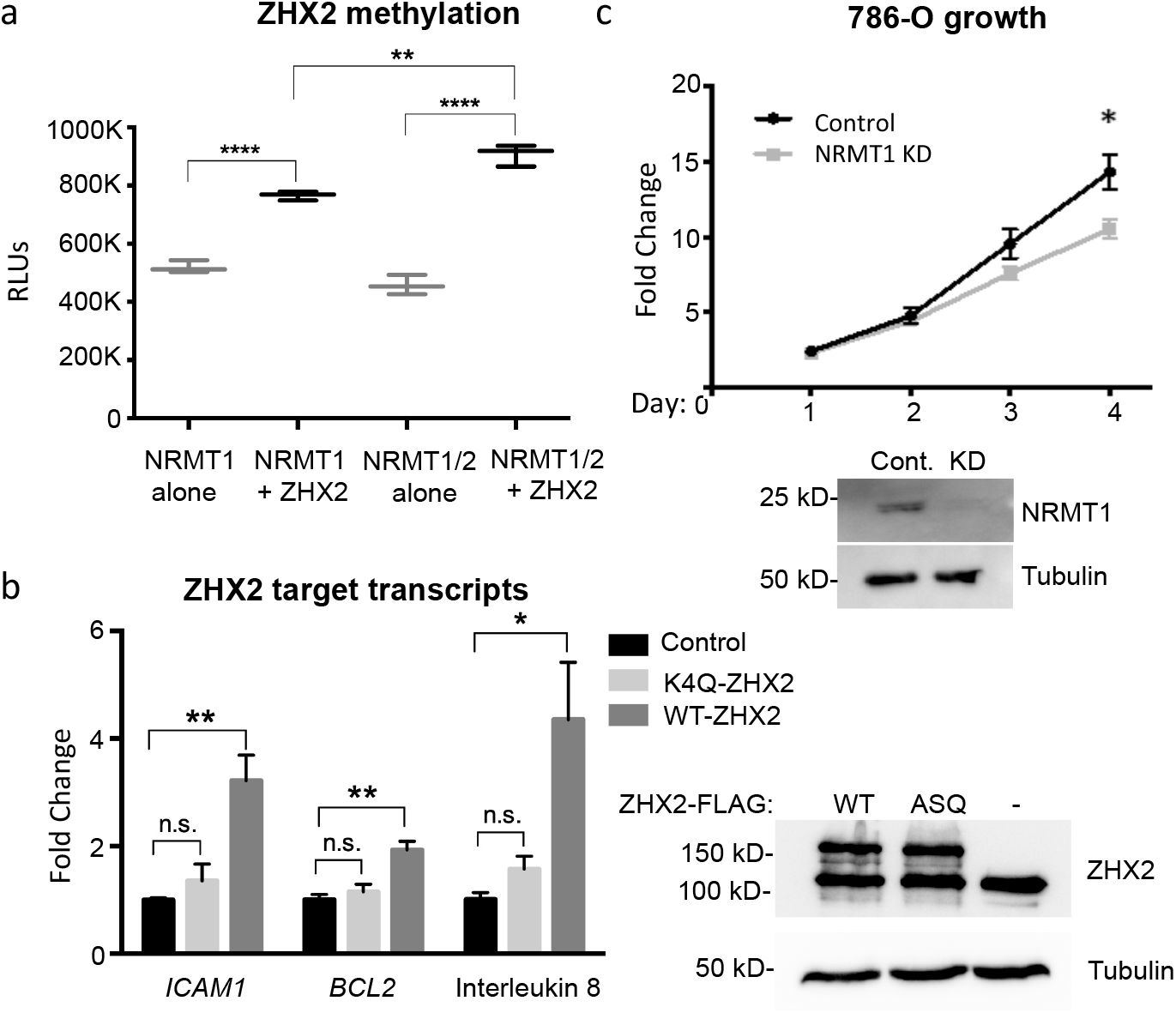
ZHX2 is methylated by NRMT1 and Nα-methylation regulates ZHX2 function. (a) *In vitro* methyltransferase assays show ZHX2 peptide is methylated by recombinant NRMT1. Addition of NRMT2 significantly increases the methylation activity above NRMT1 alone. (b - left) qRT-PCR analysis of the ZHX2 target transcripts *ICAM1, BCL2*, and interleukin 8 in untransfected, K4Q-ZHX2-GFP overexpressing, or WT-ZHX2-GFP overexpressing 786-O cells shows a significant increase in expression of all three genes when comparing WT overexpression to untransfected. K4Q overexpression does not significantly alter transcript expression as compared to untransfected. (b – right) Western blot confirms equal expression of WT and K4Q-ZHX2-GFP. Tubulin is used as a loading control. (c - upper) Growth curves of control and NRMT1 knockdown (KD) 786-O cells show NRMT1 knockdown can significantly slow growth after 4 days. (c - lower) Western blot confirming knockdown of NRMT1 in 786-O cells; tubulin is used as a loading control. * denotes p < 0.05, ** denotes p < 0.01, and **** denotes p < 0.0001 as determined by unpaired t-test. n = 3. Error bars represent ± standard error of the mean (SEM).

We have also previously shown loss of Nα-methylation of another non-canonical target, MYL9, impairs its activity as a transcriptional activator (10). Mutation of lysine 4 in the consensus sequence of MYL9 to a glutamine (K4Q) inhibits the binding of MYL9 to target promoter regions and decreases their expression levels (10). To test if Nα-methylation of ZHX2 also affects its ability to function as a transcription factor, we made a similar K4Q mutant of ZHX2. Besides regulation of *Mup* and *Cyp* genes in the liver, ZHX2 also regulates expression of NF-κB targets genes during clear cell renal cell carcinoma (ccRCC) progression (24). WT ZHX2-GFP or methyl-deficient K4Q-ZHX2-GFP were overexpressed in 786-O ccRCC cells and expression of the NF-κB targets, *BCL2, IL8*, and *ICAM1* were quantified using qRT-PCR. Overexpression of WT ZHX2-GFP resulted in increased expression of all three transcripts as compared to untransfected controls (Figure 5(b)). However, overexpression of K4Q-ZHX2-GFP resulted in a significantly lower expression of all three transcripts as compared to WT ZHX2-GFP expression, which was also not significantly different from untransfected controls (Figure 5(b)). Western blotting confirmed altered ZHX2 target expression was not due to unequal expression of the WT and K4Q-ZHX2 proteins (Figure 5(b)). These data support the RNA-seq data and indicate Nα-methylation regulates the transcription factor activity of ZHX2.

As ZHX2 controls oncogenic NF-κB signaling in ccRCC, its loss results in decreased 786-O proliferation (24). To determine if loss of NRMT1 results in a similar decrease in growth, we used a lentivirus expressing an shRNA against human NRMT1 to knockdown NRMT1 expression in 786-O cells (Figure 5(c)) and performed cell proliferation assays. Human 786-O cells transduced with lentivirus expressing an shRNA against mouse NRMT1 were used as control cells (Figure 5(c)). By the fourth day, it was evident that NRMT1 knockdown significantly slowed 786-O growth (Figure 5(c)). These data show that not only does Nα-methylation regulate ZHX2 function, but loss of NRMT1 can phenocopy a ZHX2 loss of function phenotype.

### ZHX2 function and liver degeneration

While down-regulation of *Mup* expression and overexpression of *Cyp* expression have both been shown to disrupt liver metabolic function (35–37) and are likely contributing to the observed liver degenerative phenotypes seen in *Nrmt1^-/-^* mice, ZHX2 also plays a role in liver development through its postnatal repression of the liver fetal development regulators AFP, GPC3, and H19 (19, 20). All three are expressed at high levels in the fetal liver but are subsequently silenced during postnatal development by ZHX2 (19, 20). AFP and GPC3 also play a postnatal role in liver stem cell (oval cell) regulation, and abnormal postnatal H19 expression promotes liver fibrosis (38–41). To determine if early *Afp, Gpc3*, or *H19* repression was altered in *Nrmt1^-/-^* mice, we performed qRT-PCR analysis on WT and *Nrmt1^-/-^* livers at birth (P0) and 14 days after birth (P14). While we see no significant change in *Afp* silencing between P0 and P14 (Figure 6(a)), *Gpc3* and *H19* expression is significantly de-repressed in *Nrmt1^-/-^* mice by 14 days (Figure 6(b,c)). These data indicate the transcription factor activity of ZHX2 is altered in *Nrmt1^-/-^* mice in a substrate-specific manner and suggest, in addition to misregulation of liver metabolic pathways, loss of ZHX2 activity contributes to the liver phenotype seen in *Nrmt1^-/-^* mice by altering developmental pathways as well.

**Figure 6.**
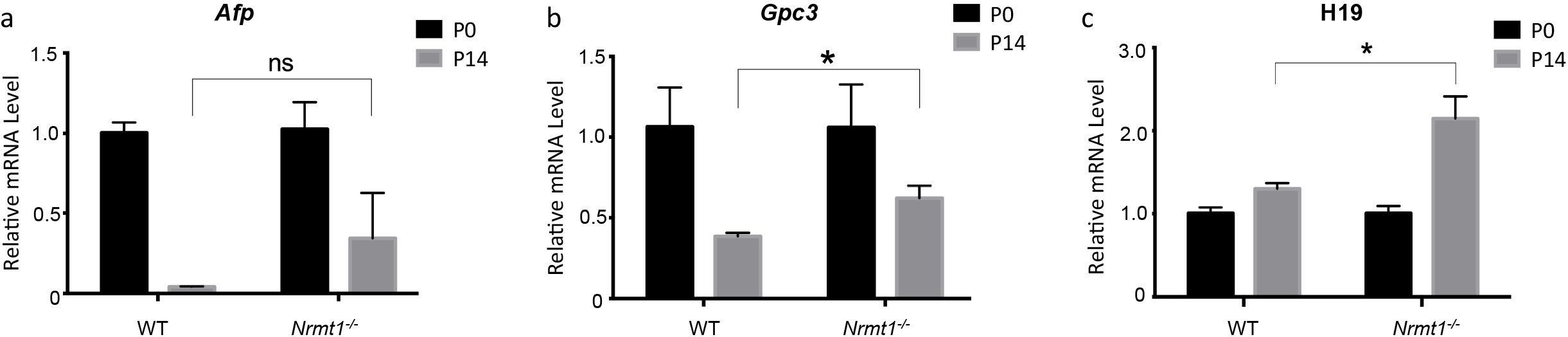
Postnatal *Gpc3* and H19 expression is de-repressed in *Nrmt1^-/-^* livers. qRT-PCR analysis of wild-type (WT) and Nrmt1 knockout (*Nrmt1^-/-^*) mouse livers reveals (a) *Afp* expression is not de-repressed in *Nrmt1^-/-^* livers between 0 and 14 days after birth. However, (b) *Gpc3* and (c) *H19* expression is significantly de-repressed by 14 days. * p < 0.05 as determined by unpaired t-test. n = 3. Error bars represent ± standard error of the mean (SEM).

## Discussion

The numerous phenotypes exhibited by *Nrmt1^-/-^* mice, including premature hair loss, cystic ovaries, and liver degeneration (12), speak to the wide variety of targets of Nα-methylation that are misregulated when NRMT1 is lost. Here we characterize ZHX2 as a novel substrate of NRMT1 and demonstrate how the loss of Nα-methylation disrupts the ability of ZHX2 to regulate its targets in the mouse liver, including *Mups, Cyps, Gpc3*, and *H19*, and in renal cancer cells, including NF-κB targets.

Though we found many transcripts misregulated in the *Nrmt1^-/-^* livers, we predict those regulated by ZHX2 are a driving force in the observed liver phenotypes, as BALB/cJ mice, which harbor a hypomorphic ZHX2 mutation, are also highly susceptible to liver damage (42). Exactly what downstream ZHX2 targets are involved in promoting liver degeneration remains to be elucidated, but our data suggest both metabolic and developmental pathways play a role. Increased *Cyp* expression has been shown to promote oxidative stress in the liver, and decreased *Mup1* expression alters both glucose and lipid metabolism (35–37). Postnatal *Gpc3* expression is normally isolated to hepatic progenitor/oval cells (38), so its de-repression could abnormally expand the postnatal oval cell population. Abnormal postnatal *H19* expression promotes liver fibrosis, in part, through inhibition of hepatic zinc finger E-box binding homeobox 1 (ZEB1), which is then unable to repress expression of epithelial cell adhesion molecule (EpCAM) (39). H19 has also been shown to act as a sponge for miR-148a, which results in stabilized TGF-β receptor I and the promotion of epithelial-mesenchymal transition (EMT) (40). While our data suggest many ZHX2 downstream targets contribute to the observed phenotypes, we saw no difference in *Afp* postnatal de-repression, indicating Nα-methylation does not control ZHX2 transcription factor activity at all sites. We predict the effect of Nα-methylation may be DNA sequence dependent or reliant on whether ZHX2 homodimerizes, heterodimerizes with ZHX1/ZHX3, or is bound to alternate complex members including NF-YA (15, 16).

The liver is likely not the only organ in *Nrmt1^-/-^* mice where ZHX2 activity is affecting phenotypic outcomes. We have recently shown that the two neural stem cell (NSC) niches in the brain, the subventricular zone (SVZ) of the lateral ventricles and the subgranular zone (SGZ) of the dentate gyrus, exhibit postnatal misregulation of the cell cycle (43). The NSCs from these niches show precocious proliferation followed by premature depletion of the quiescent stem cell pool (43). This results in both neurodegenerative phenotypes and cognitive impairments in *Nrmt1^-/-^* mice (43). Through its interactions with ephrin-B1, ZHX2 has been shown to regulate neural progenitor cell maintenance and prevent premature neuronal proliferation (44), indicating it may also be an important downstream target of NRMT1 in the brain. Similar RNA-sequencing analysis of cells in the SVZ and SGZ of *Nrmt1^-/-^* mice will help further identify ZHX2 targets regulated by Nα-methylation in the brain. It will also help elucidate tissuespecific signaling pathways regulated by ZHX2.

Besides Nα-methylation, the N-terminal sequence of ZHX2 (ASK) allows it to be targeted for Nα-acetylation (5). Our lab previously characterized MYL9 as the first protein regulated by both Nα-methylation and Nα-acetylation (10). While Nα-methylation regulated the nuclear roles of MYL9 as a transcription factor, Nα-acetylation regulated the cytoplasmic roles of MYL9 in cell migration (10). We have shown here that the nuclear role of ZHX2 as a transcription factor is also regulated by Nα-methylation, but ZHX2 is also found in the cytoplasm. Though there are currently no known cytoplasmic roles of ZHX2, its cytoplasmic localization does increase in disease states, including during hepatocellular carcinoma progression (22). Future studies will be aimed at identifying novel cytoplasmic functions of ZHX2 and determining if they are regulated by Nα-acetylation.

The thirteen amino-terminal acids of ZHX2 are incredibly highly conserved, implying there is a functional relevance for the N-terminus (45). Thus, regulation of ZHX2 function through Nα-methylation, which is more rapid than upregulating production of more ZHX2 protein, is a sensitive method for modulating the expression of ZHX2 targets. Both in the mouse liver and in renal cancer cells, loss of Nα-methylation is sufficient to disrupt the activity of ZHX2 as a transcription factor, and Nα-acetylation may be providing a separate regulatory role as well. Continued characterization of how Nα-PTMs affect ZHX2 function will give us a better understanding of how ZHX2 regulates both developmental and oncogenic processes.

## Acknowledgments

We thank Meghan Heaney and Lindsay Bonsignore for their contributions to the manuscript and Elizabeth Hudson at the University of Louisville Center for Genetics and Molecular Medicine (CGeMM) for performing the RNA sequencing. This work was supported by a Next-Gen Pilot grant from CGeMM and research grant from the National Institutes of Health to C.E.S.T [GM112721].

---

**Supplemental Table 1.**
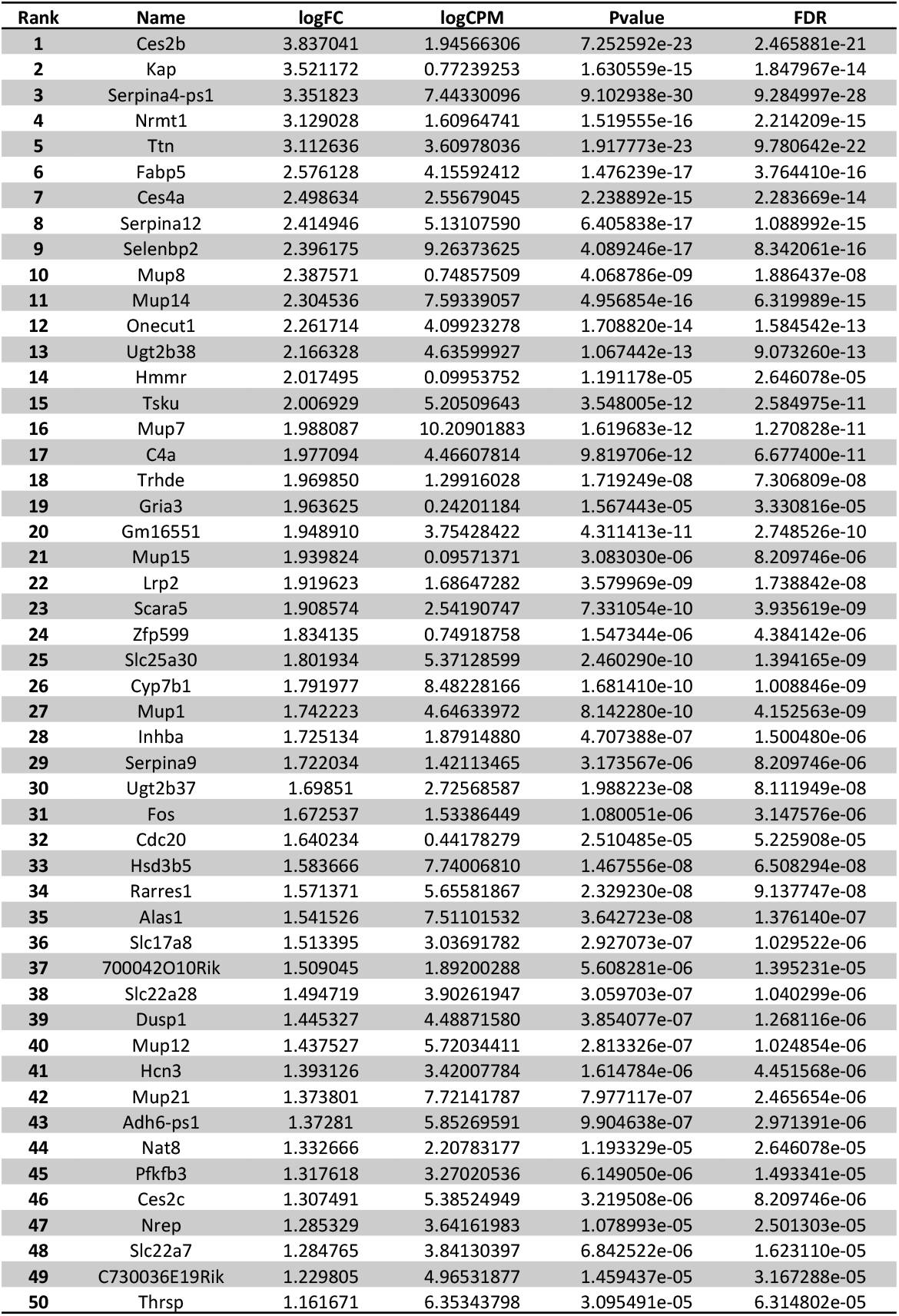
Top 50 transcripts down-regulated in *Nrmt1^-/-^* mice.

| Rank | Name | logFC | logCPM | Pvalue | FDR |
| --- | --- | --- | --- | --- | --- |
| 1 | Ces2b | 3.837041 | 1.94566306 | 7.252592e-23 | 2.465881e-21 |
| 2 | Kap | 3.521172 | 0.77239253 | 1.630559e-15 | 1.847967e-14 |
| 3 | Serpina4-ps1 | 3.351823 | 7.44330096 | 9.102938e-30 | 9.284997e-28 |
| 4 | Nrmt1 | 3.129028 | 1.60964741 | 1.519555e-16 | 2.214209e-15 |
| 5 | Ttn | 3.112636 | 3.60978036 | 1.917773e-23 | 9.780642e-22 |
| 6 | Fabp5 | 2.576128 | 4.15592412 | 1.476239e-17 | 3.764410e-16 |
| 7 | Ces4a | 2.498634 | 2.55679045 | 2.238892e-15 | 2.283669e-14 |
| 8 | Serpina12 | 2.414946 | 5.13107590 | 6.405838e-17 | 1.088992e-15 |
| 9 | Selenbp2 | 2.396175 | 9.26373625 | 4.089246e-17 | 8.342061e-16 |
| 10 | Mup8 | 2.387571 | 0.74857509 | 4.068786e-09 | 1.886437e-08 |
| 11 | Mup14 | 2.304536 | 7.59339057 | 4.956854e-16 | 6.319989e-15 |
| 12 | Onecut1 | 2.261714 | 4.09923278 | 1.708820e-14 | 1.584542e-13 |
| 13 | Ugt2b38 | 2.166328 | 4.63599927 | 1.067442e-13 | 9.073260e-13 |
| 14 | Hmmr | 2.017495 | 0.09953752 | 1.191178e-05 | 2.646078e-05 |
| 15 | Tsku | 2.006929 | 5.20509643 | 3.548005e-12 | 2.584975e-11 |
| 16 | Mup7 | 1.988087 | 10.20901883 | 1.619683e-12 | 1.270828e-11 |
| 17 | C4a | 1.977094 | 4.46607814 | 9.819706e-12 | 6.677400e-11 |
| 18 | Trhde | 1.969850 | 1.29916028 | 1.719249e-08 | 7.306809e-08 |
| 19 | Gria3 | 1.963625 | 0.24201184 | 1.567443e-05 | 3.330816e-05 |
| 20 | Gm16551 | 1.948910 | 3.75428422 | 4.311413e-11 | 2.748526e-10 |
| 21 | Mup15 | 1.939824 | 0.09571371 | 3.083030e-06 | 8.209746e-06 |
| 22 | Lrp2 | 1.919623 | 1.68647282 | 3.579969e-09 | 1.738842e-08 |
| 23 | Scara5 | 1.908574 | 2.54190747 | 7.331054e-10 | 3.935619e-09 |
| 24 | Zfp599 | 1.834135 | 0.74918758 | 1.547344e-06 | 4.384142e-06 |
| 25 | Slc25a30 | 1.801934 | 5.37128599 | 2.460290e-10 | 1.394165e-09 |
| 26 | Cyp7b1 | 1.791977 | 8.48228166 | 1.681410e-10 | 1.008846e-09 |
| 27 | Mup1 | 1.742223 | 4.64633972 | 8.142280e-10 | 4.152563e-09 |
| 28 | Inhba | 1.725134 | 1.87914880 | 4.707388e-07 | 1.500480e-06 |
| 29 | Serpina9 | 1.722034 | 1.42113465 | 3.173567e-06 | 8.209746e-06 |
| 30 | Ugt2b37 | 1.69851 | 2.72568587 | 1.988223e-08 | 8.111949e-08 |
| 31 | Fos | 1.672537 | 1.53386449 | 1.080051e-06 | 3.147576e-06 |
| 32 | Cdc20 | 1.640234 | 0.44178279 | 2.510485e-05 | 5.225908e-05 |
| 33 | Hsd3b5 | 1.583666 | 7.74006810 | 1.467556e-08 | 6.508294e-08 |
| 34 | Rarres1 | 1.571371 | 5.65581867 | 2.329230e-08 | 9.137747e-08 |
| 35 | Alas1 | 1.541526 | 7.51101532 | 3.642723e-08 | 1.376140e-07 |
| 36 | Slc17a8 | 1.513395 | 3.03691782 | 2.927073e-07 | 1.029522e-06 |
| 37 | 700042O10Rik | 1.509045 | 1.89200288 | 5.608281e-06 | 1.395231e-05 |
| 38 | Slc22a28 | 1.494719 | 3.90261947 | 3.059703e-07 | 1.040299e-06 |
| 39 | Dusp1 | 1.445327 | 4.48871580 | 3.854077e-07 | 1.268116e-06 |
| 40 | Mup12 | 1.437527 | 5.72034411 | 2.813326e-07 | 1.024854e-06 |
| 41 | Hcn3 | 1.393126 | 3.42007784 | 1.614784e-06 | 4.451568e-06 |
| 42 | Mup21 | 1.373801 | 7.72141787 | 7.977117e-07 | 2.465654e-06 |
| 43 | Adh6-ps1 | 1.37281 | 5.85269591 | 9.904638e-07 | 2.971391e-06 |
| 44 | Nat8 | 1.332666 | 2.20783177 | 1.193329e-05 | 2.646078e-05 |
| 45 | Pfkfb3 | 1.317618 | 3.27020536 | 6.149050e-06 | 1.493341e-05 |
| 46 | Ces2c | 1.307491 | 5.38524949 | 3.219508e-06 | 8.209746e-06 |
| 47 | Nrep | 1.285329 | 3.64161983 | 1.078993e-05 | 2.501303e-05 |
| 48 | Slc22a7 | 1.284765 | 3.84130397 | 6.842522e-06 | 1.623110e-05 |
| 49 | C730036E19Rik | 1.229805 | 4.96531877 | 1.459437e-05 | 3.167288e-05 |
| 50 | Thrsp | 1.161671 | 6.35343798 | 3.095491e-05 | 6.314802e-05 |

**Supplemental Table 2.**
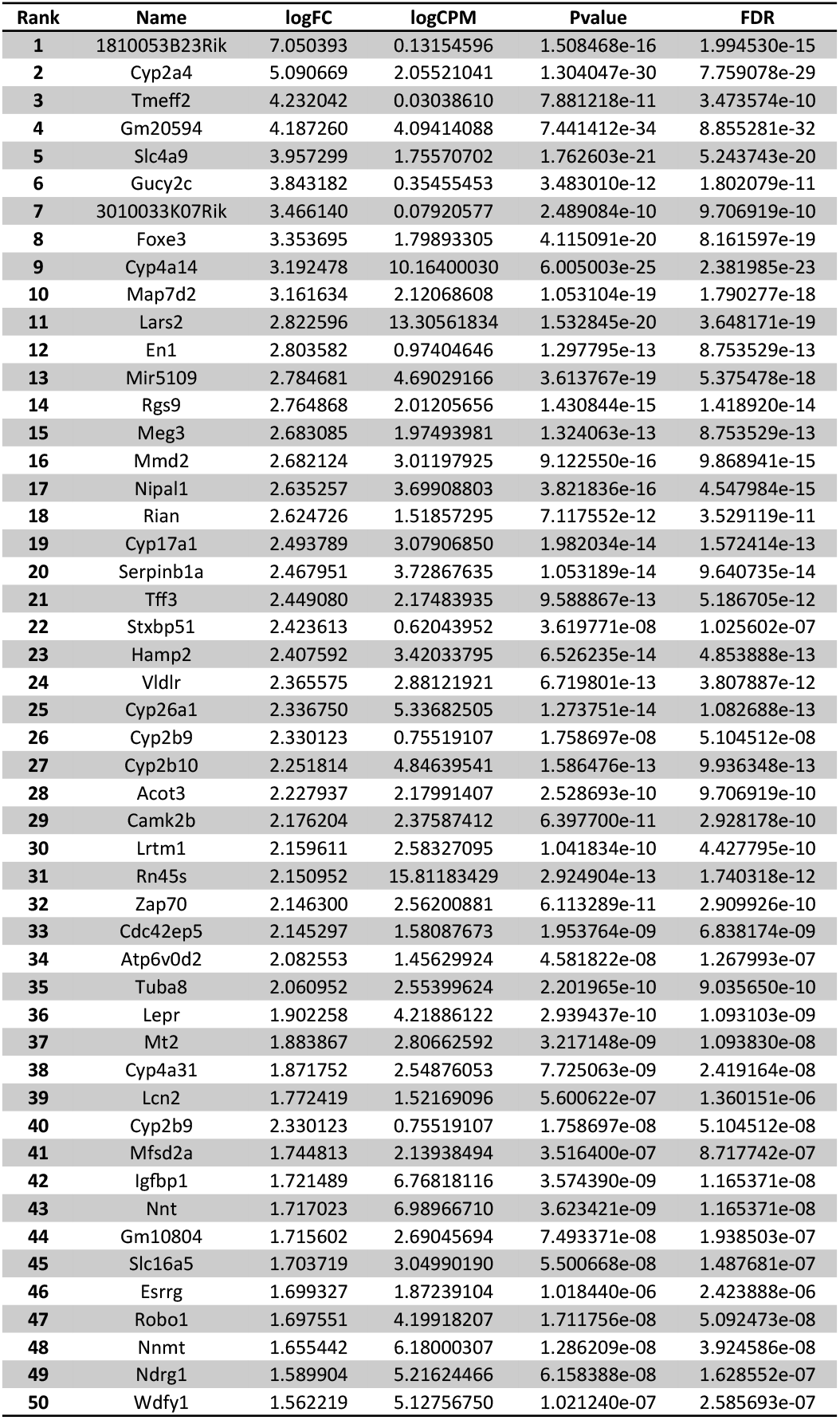
Top 50 transcripts upregulated in *Nrmt1^-/-^* mice.

